# A novel role for cortical acetylcholine in object memory updating

**DOI:** 10.1101/2020.02.08.940064

**Authors:** Kristen H. Jardine, Cassidy E. Wideman, Chelsea MacGregor, Cassandra Sgarbossa, Dean Orr, Krista A. Mitchnick, Boyer D. Winters

**Affiliations:** Department of Psychology and Collaborative Neuroscience Program, University of Guelph, Guelph, ON, Canada, N1G 2W1

**Author notes:** Authors contributed equally to this work (Kristen H. Jardine, Cassidy E. Wideman). Corresponding Author: B.D. Winters, Department of Psychology, University of Guelph, 50 Stone Rd. East, Guelph, ON Canada, N1G2W1.

## Abstract

Reactivated long-term memories can become labile and sensitive to modification. Memories in this destabilized state can be weakened or strengthened, but there is limited research characterizing the mechanisms underlying retrieval-induced qualitative updates (i.e., information integration). We have previously implicated cholinergic transmission in object memory destabilization. Here we present a novel rodent paradigm developed to assess the role of this cholinergic mechanism in qualitative memory updating. The post-reactivation object memory modification (PROMM) task exposes rats to contextual information following object memory reactivation. Subsequent object exploratory performance suggests that the contextual information is integrated with the original memory in a reactivation- and time-dependent manner. This effect is blocked by interference with M_1_ muscarinic receptors and several downstream signals in perirhinal cortex. These findings therefore demonstrate a hitherto unacknowledged cognitive function for acetylcholine with important implications for understanding the dynamic nature of long-term memory storage in the normal and aging brain.

## Introduction

When first acquired, memories exist in a labile state and require protein synthesis-dependent consolidation to stabilize for long-term storage^1,2^. Following the presentation of reminder cues, however, consolidated long-term memories can be destabilized and again rendered labile, necessitating a second protein synthesis-dependent re-stabilization process referred to as reconsolidation^3–5^. The process of reconsolidation has been widely posited to play a role in adaptive memory updating, enabling the maintenance of memory accuracy and relevance over time^6,7^. Indeed, studies in rodents and humans have demonstrated post-reactivation modification by manipulating the strength of fear memory, providing evidence for memory trace weakening or erasure^4,8^, as well as strengthening through targeted additional training ^9^. However, these studies do not directly assess the integration of new, relevant information presented during the reconsolidation window to update the content of established long-term memories. The distinction between this ‘qualitative’ memory updating and the previously demonstrated ‘quantitative’ changes may seem subtle, but it is likely significant when considering the dynamic nature of long-term storage for different types of material.

As informative as past studies have been, fear conditioning^9–14^ or other conditioning-based learning (e.g., drug associated cues^15^) bear little resemblance to human declarative memory, which exerts significant influence over day-to-day behaviour and is subject to regular qualitative modification. Indeed, the modifiability of human declarative memory – beyond merely strengthening and weakening - has long been acknowledged^16–19^. Perhaps most convincing is the occurrence of false memories, whereby incorrect information integrates into long-term memory and persists over time^20^. The phenomenon of reconsolidation may provide a link between understanding the cognitive basis of memory modification and its underlying neurobiology^21^. In order to bridge the gap between these fields, a feasible animal model of declarative memory modification that provides evidence for the incorporation of new information into an existing memory is required.

Object recognition tests have been used extensively to characterize animal models of amnesia^22,23^ and are widely accepted as standard procedures for studying declarative-like memory in the rodent literature^23^. Previous studies have adapted spontaneous object recognition tasks to provide evidence for memory updating through reconsolidation mechanisms; typically, at the time of reactivation, the reminder cue is presented with updating information, such as a new object or object location^25–27^. While these studies have made valuable contributions, the structure of these tasks, whereby the reactivation episode is combined with the presentation of updating information, presents two problems. First, this experience could be encoded as a completely novel learning episode rather than an opportunity to update the original memory; and second, it prevents assessment of the potential for memory modification within the post-reactivation reconsolidation window. A cleaner procedure for studying the neural bases of memory updating would present the updating information following memory reactivation during the labile window. One of the most convincing demonstrations of reconsolidation-mediated constructive memory updating in humans utilized a similar methodology. In this study, information about the contents of a new list of objects was incorporated into the memory of a list memorized earlier, but only if the second list was presented shortly after reactivation of the memory for the first list. Thus, the updating effect was reactivation- and time-dependent, as well as a constructive process^28^.

Here we present a novel paradigm for use with rodents to study the behavioural and neural mechanisms of reconsolidation-based memory updating. In this post-reactivation object memory modification (PROMM) task, new contextual information appears to be incorporated into a previously acquired object memory when presented while the object memory trace is labile following reactivation. We also use the PROMM task to investigate the neurobiological mechanisms underlying object memory updating. Previously, our group demonstrated the requirement of cholinergic signaling at M_1_ muscarinic receptors (mAChRs) in perirhinal cortex (PRh) for destabilization of object memories^29,30^. This effect appears to be mediated by signalling downstream of the M_1_ receptor, including the second messenger inositol triphosphate (IP_3_) and activation of the ubiquitin proteasome system (UPS), which is involved in degrading synaptic proteins and is thought to underlie destabilization at the synaptic level^14,30^. We hypothesized that this mechanism of memory destabilization is necessary for reconsolidation-mediated memory updating, but until now there has been no viable rodent model to study this question. Using the PROMM task, the current study presents evidence in support of this hypothesis, implicating cortical acetylcholine (ACh) in a cognitive function that is subtly, but clearly, distinct from its established role in new learning.

## Results

### Retrieval enables constructive and time-dependent object memory updating in rats

In the PROMM task, rats explore an object during a sample phase, and the object memory is reactivated 24h later with a brief re-exposure to the identical training objects and context. After the object memory is reactivated, and consequently destabilized, the rat is immediately placed into an empty alternate context (see Methods; Fig. 1). Here, we predict that the alternate contextual information will incorporate into the labile object memory representation. On test day, an additional 24h later, rats explore the sampled objects in either the same alternate context as seen post-reactivation, or a different alternate context. If the alternate context presented post-reactivation successfully integrates with the reactivated object memory, rats should behave as if this test phase object-context combination is familiar, even though the objects have never been directly experienced in that context. When the objects are presented in combination with an alternate context during test phase that is *different* than the context explored post-reactivation, rats should behave as if this object-context configuration is unfamiliar, and therefore demonstrate novelty-induced increases in object exploration^31^. As predicted, rats showed significantly lower total object exploration in the same alternate context condition compared to the different alternate context condition, *t*(11)=−3.29, *p*=.007 (Fig. 2a).

**Figure 1.**
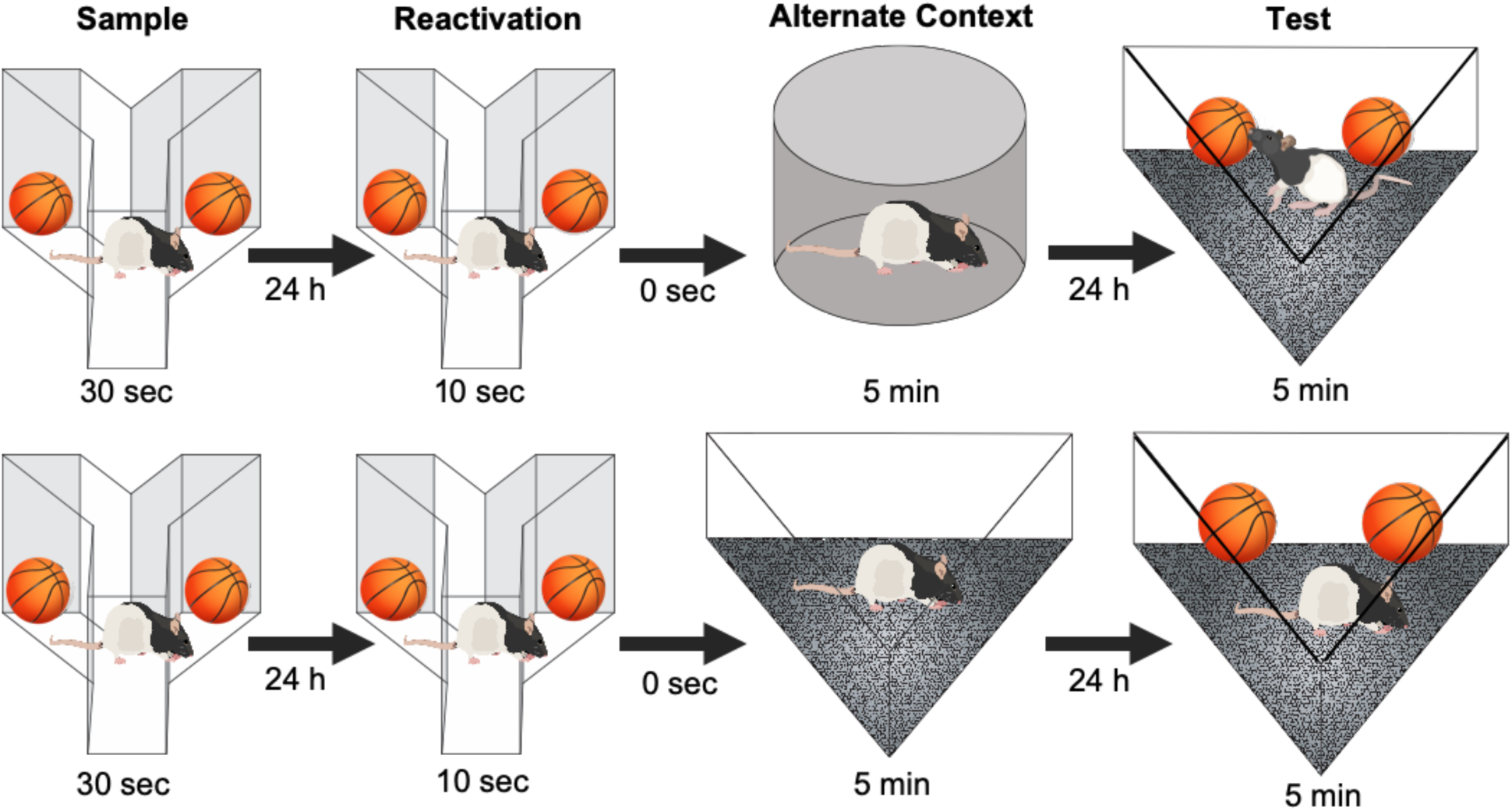
Illustration of the post-reactivation object memory modification (PROMM) task. The rat samples a pair of identical objects in a Y-apparatus, and the object memory is reactivated 24 h later with a brief re-exposure to the same objects in the same Y-apparatus. After memory reactivation in the Y-apparatus, the rat is immediately placed into an empty alternate context to which it has previously been habituated, but where it has never explored objects. In the test phase, 24 h after memory reactivation, the rat is given 5 min to explore the sample objects in one of two alternate contexts: either the alternate context explored immediately after reactivation (‘same alternate context’ condition; bottom of figure) or another previously habituated alternate context not presented on that trial (‘different alternate context’ condition; top of figure).

**Figure 2.**
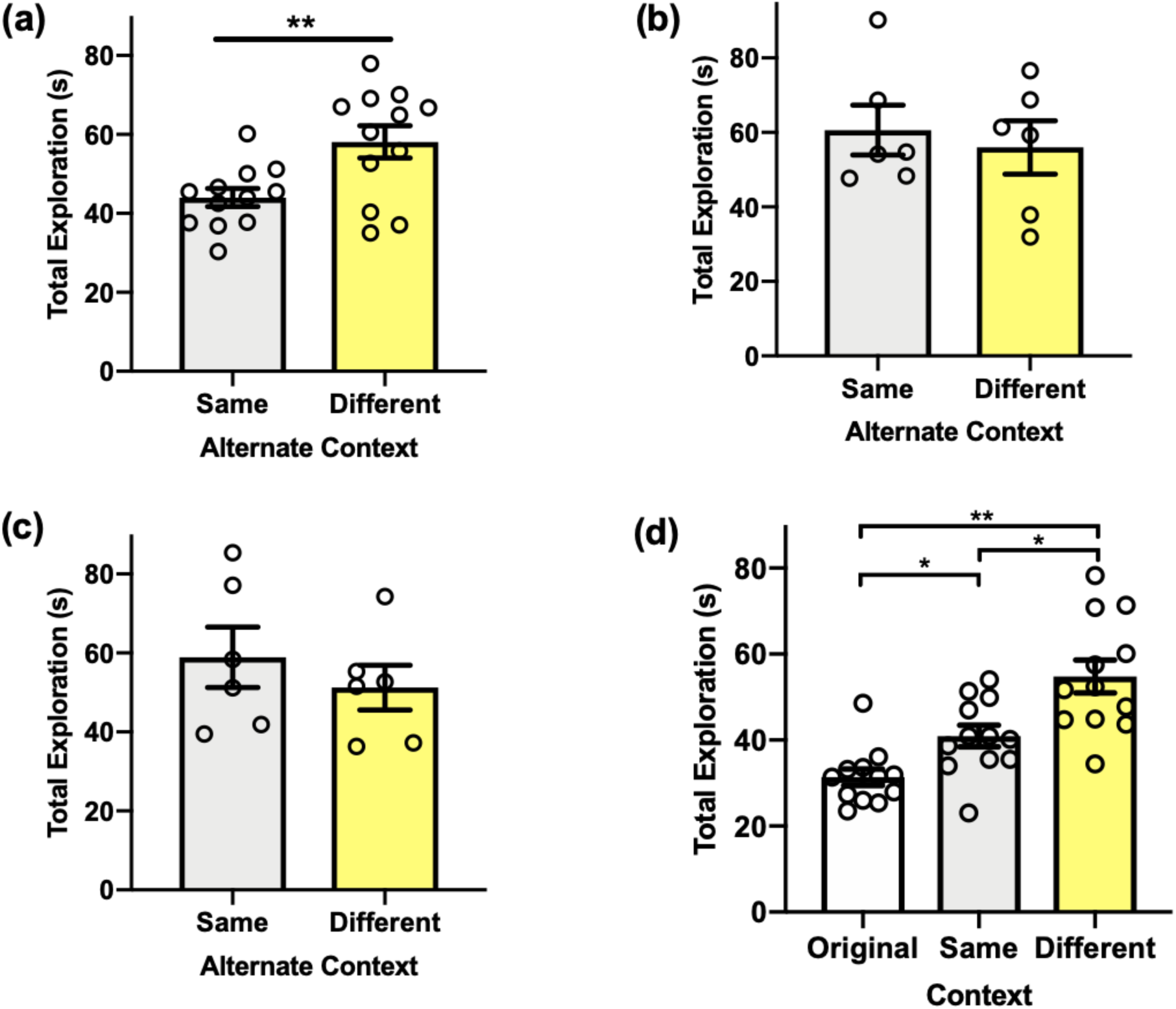
Behavioural performance in the PROMM task. **(a)** Total exploration of the sampled objects in the test phase is greater when the objects are presented in a different apparatus than the one explored immediately after memory reactivation (n=12). This pattern is consistent with the interpretation that rats perceive the same alternate context-object configuration as more familiar than the different alternate context-object configuration. **(b)** When the reactivation phase is omitted, there is no difference between test phase exploration in the same alternate context and different alternate context conditions (n=12). **(c)** There is no difference in test phase object exploration between the alternate context conditions when the alternate arena is presented 6 h after object memory trace reactivation (n=12). **(d)** Memory for the original object-context configuration appears to be intact when exploration of the sample objects is measured in the Y-apparatus (i.e., the original sample context) during the test phase (n=12). Object exploration in the Y-apparatus during test is lower than in either the different alternate context or the same alternate context, indicating that the original object memory is not detrimentally affected by post-reactivation updating. Object exploration in the same alternate context condition was lower than object exploration in the different alternate context, replicating the effect reported in (a). \**p*<.05, \*\**p*<.01.

Next, we looked to confirm that this apparent object memory updating is indeed reliant on memory trace reactivation. To this end, we again ran the PROMM task, but with the reactivation phase omitted. In the absence of explicit object memory reactivation, there was no statistical difference between object exploration in the same alternate context condition and the different alternate context condition, *t*(10)=.476, *p*=.645 (Fig. 2b).

It is generally accepted that memory modification must occur within a distinct window of time following reactivation, before the memory trace is reconsolidated and no longer modifiable^5^. To assess the time-dependency of the updating effect in the PROMM task, we ran the task as above but with the empty alternate context presented 6h post-reactivation, presumably outside of the reconsolidation window. An independent samples t-test revealed that postponing the presentation of the alternate contextual information abolished the behavioural effect, and there was no statistical difference between test phase object exploration in the same and different alternate context conditions, *t*(10)=.806, *p*=.439 (Fig. 2c).

A key characteristic of the PROMM task is its demonstration of reactivation-based updating, as opposed to reactivation-induced “erasure”. One objective of the PROMM task is to update the original memory with new information while keeping the original memory intact. We therefore aimed to verify that the original object memory established in the sample phase of the PROMM task could still be successfully retrieved after the object memory was modified following the reactivation phase. To investigate this, we measured exploration of the objects on test day in the Y-apparatus, the original context in which the objects were sampled. A repeated measures one-way analysis of variance (ANOVA) revealed a significant effect of context, *F*(2,22)=19.840, p<.001 (Fig. 2d). As previously shown, exploration of the sampled objects in the same alternate context was significantly lower than exploration of the sampled objects in the different alternate context, *t*(11)= -2.927, *p*=.014. Moreover, total exploration in the original sample context was significantly lower than exploration in both the same alternate context, *t*(11)= -4.484, *p*=.001, and the different alternate context, *t*(11)= -5.930, *p*<.001. Thus, the object-context configuration including the original context from the sample phase of the task (i.e., the Y-apparatus) was treated as the most familiar; this is perhaps not surprising because the rats are exposed to this configuration explicitly during both the sample phase and the reactivation phase.

### Object memory updating in the PROMM task requires proteasome activity

We next aimed to confirm the necessity of UPS activity for the observed memory updating effect. As mentioned, proteasomes of the UPS are suspected to facilitate the synaptic degradation that it posited to underlie memory destabilization^14^, and we have previously implicated UPS activity in object memory labilization^30^. In order to verify that UPS activation is similarly important for the object memory updating measured by the PROMM task, we evaluated the effect of blocking 26S proteasome activity prior to memory reactivation directly within PRh using pre-reactivation intra-PRh microinjections of the proteasome inhibitor clasto-lactacystin β-lactone (β-lac). This prevented the reduction in object exploration typically seen in the same alternate context condition on test day (Fig. 3). A 2×2 factorial ANOVA revealed a significant interaction between the alternate context conditions (same, different) and drug conditions (vehicle, β-lac), *F*(1,49)=7.025, *p*=.011. Further, a main effect of drug was revealed, *F*(1,49)=4.996, *p*=.030. Importantly, rats in the vehicle/same alternate context group had significantly lower test phase object exploration compared to subjects in the β-lac/same alternate context condition, *t*(20.148)= −3.340, *p*=.003. Rats in the vehicle/same alternate context condition also had lower exploration than rats in the vehicle/different alternate context condition, *t*(24)= -3.337, *p*=.003, and β-lac/different alternate context condition, *t*(25)= -2.775, *p*=.010.

**Figure 3.**
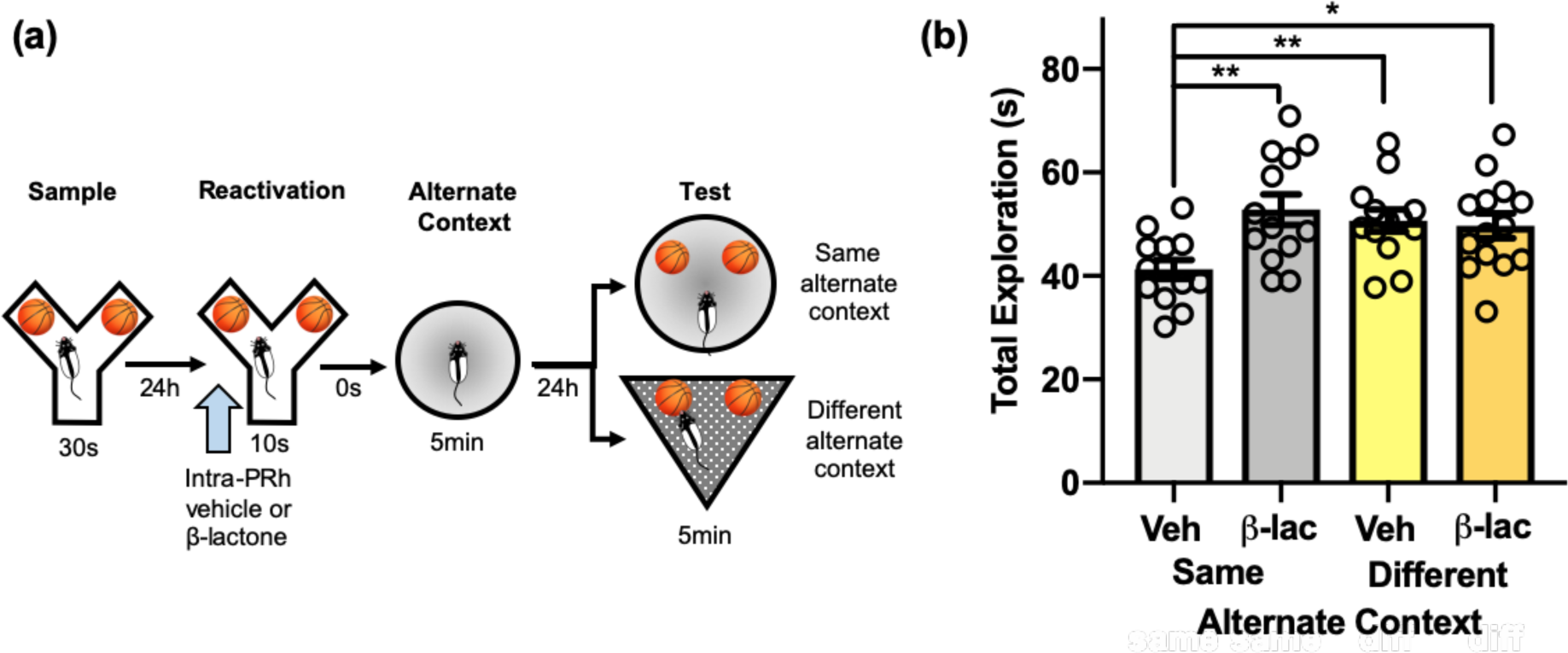
Proteasome inhibition in PRh prevents object memory updating in the PROMM task. **(a)** Intracranial PRh microinjections (blue arrow) of the proteasome inhibitor β-lactone or its vehicle were administered immediately prior to object re-exposure (n=53). **(b)** Pre-reactivation β-lactone reversed the typical reduction in object exploration in the same context test condition. The (β-lactone/same alternate context condition had greater object exploration than the vehicle/same alternate context condition. Rats in the different alternate context conditions, regardless of drug, also displayed increased object exploration. \**p*<.05, \*\**p*<.01.

### Systemic mAChR antagonism prevents object memory updating in the PROMM task

We previously demonstrated that mAChR activity is necessary for object memory destabilization^29,30^. To investigate the importance of this cholinergic mechanism for reactivation-based object memory modification, we tested the effects of pre-reactivation mAChR antagonism on the apparent object memory updating observed in the PROMM task. Systemic injections of the mAChR antagonist scopolamine hydrobromide prior to the memory reactivation phase prevented the object memory updating effect (Fig. 4). A mixed measures ANOVA, with the drug conditions (vehicle, scopolamine) given within-subjects and the context conditions (same, different) implemented between-subjects, revealed a significant interaction between drug and context conditions, *F*(1,22)=5.18, *p*=.033. There was a main effect of drug, *F*(1,22)=10.09, *p*=.004, and a main effect of context, *F*(1,22)=28.12, *p*<001. In the same alternate context condition, rats administered vehicle had significantly less total object exploration in the test phase compared with rats that received scopolamine, *t*(11)=−4.04, *p*=.002. Further, rats in the vehicle/same alternate context condition also had significantly less test phase total object exploration than the vehicle/different alternate context condition, *t*(22)=−6.64, *p*<.001, and scopolamine/different alternate context condition, *t*(22)= −6.23, *p*<.001. There were no statistical interaction in the sample phase exploration, but there was a main effect of context, F(1,22)=151.87, *p*<.001. However, the different alternate context conditions had greater exploration in sample phase, and yet still explored the objects more in the test phase compared to the same alternate context groups, despite more time studying the objects during the sample phase. Also, there was a main effect of drug in reactivation exploration, F(1, 22)=12.38, p=.002. In reactivation, the group means of exploration differed by a fraction of a second, so this difference in reactivation exploration is likely not detrimental to test phase performance.

**Figure 4.**
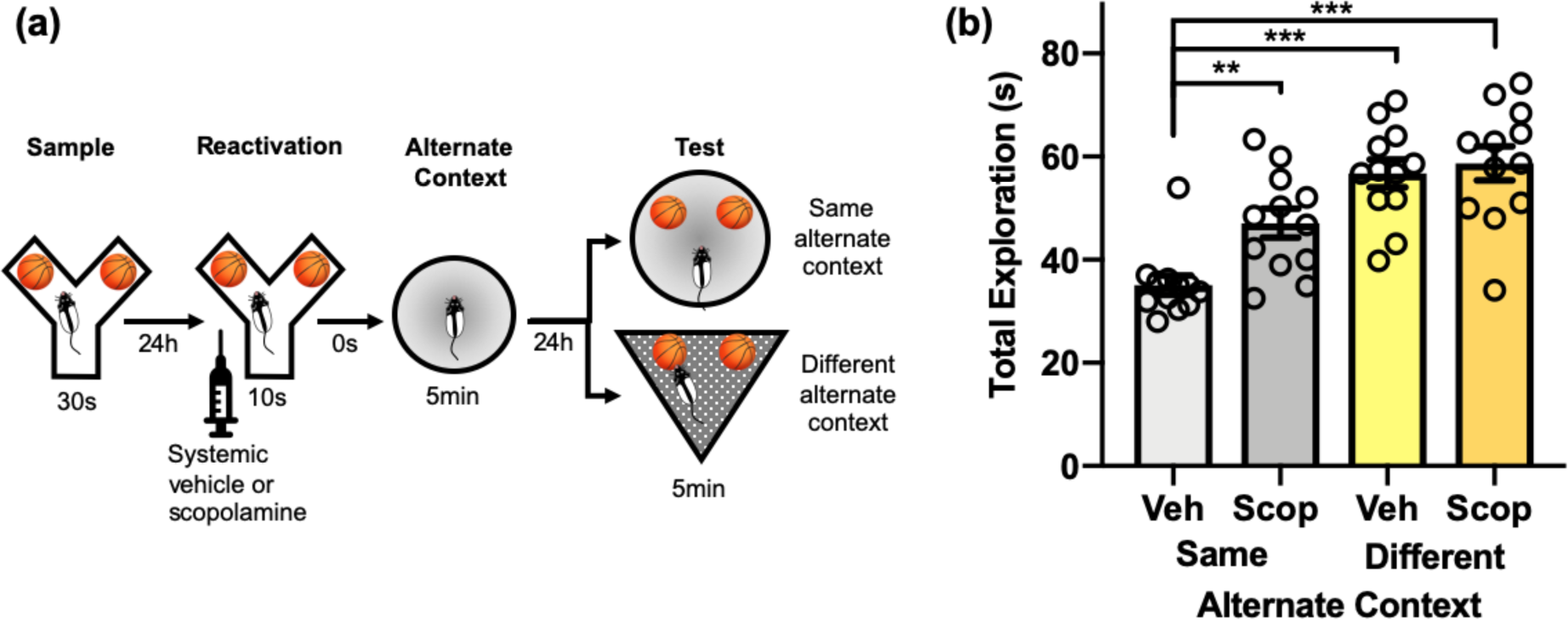
Muscarinic receptor blockade disrupts object memory updating in the PROMM task. **(a)** Intraperitoneal injections of scopolamine hydrobromide or saline were given 20 min before the reactivation phase (n=24). **(b)** Rats in the same alternate context condition that received pre-reactivation scopolamine displayed greater object exploration in the test phase, similar to the different alternate context conditions, compared to the vehicle/same alternate context condition. \*\**p*<.01, \*\*\**p*<.001

### M_1_ mAChR subtype activation in PRh is necessary for object memory updating

To further evaluate the cholinergic mechanism that facilitates constructive object memory modification, we assessed the effect of M_1_ mAChR subtype antagonism on object memory updating. Intra-PRh microinjections of pirenzepine, a selective M_1_ receptor antagonist, administered prior to the reactivation phase of the PROMM task, prevented the updating effect (Fig. 5). A repeated measures ANOVA revealed a significant interaction between drug and context conditions, *F*(1,9)=6.69, *p*=.029, as well as a main effect of drug, *F*(1,9)=25.85, *p*=.001. Similar to findings with pre-reactivation scopolamine, rats in the same alternate context condition that received microinjections of pirenzepine immediately before object memory reactivation had greater total object exploration in the test phase than rats given vehicle in the same alternate context condition, *t*(9)=−4.56, *p*=.001. Furthermore, rats in the vehicle/same alternate context condition also had lower test phase object exploration than those in the vehicle/different alternate context, *t*(9)=−3.39, *p*=.008, and pirenzepine/different alternate context, *t*(9)=−6.20, *p*<.001, conditions.

**Figure 5.**
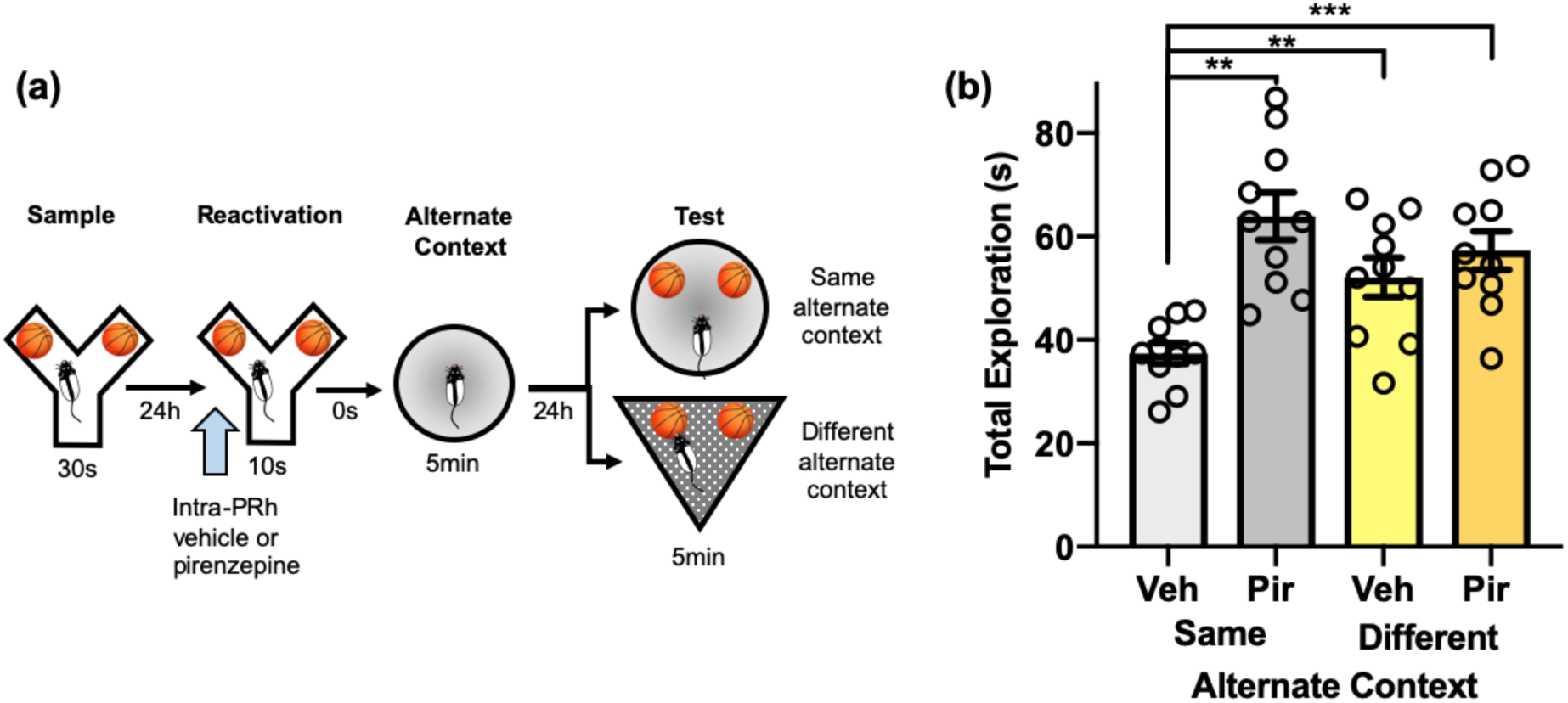
M_1_ mAChR activation within PRh is necessary for object memory updating in the PROMM task. **(a)** Intra-PRh microinjections (blue arrow) of the M_1_ mAChR antagonist pirenzepine or its vehicle were administered immediately prior to object memory reactivation in the Y-apparatus (n=10). **(b)** Test phase object exploration in the vehicle/same alternate context condition was lower than in the pirenzepine/same alternate context condition, vehicle/different alternate context condition, and pirenzepine/different alternate context condition. \*\**p*<.01, \*\*\**p*<.001

### IP_3_ receptor blockade in PRh prevents object memory updating in the PROMM task

The second messenger IP_3_, which can be stimulated by M_1_ receptor activation, binds to receptors on the endoplasmic reticulum to prompt release of intracellular Ca^2+^ stores^32^. We have previously provided evidence that this process is necessary for M_1_ receptor-induced object memory destabilization^30^, and the resultant rise in Ca^2+^ (and presumptive CaMKII mobilization) is likely a critical step in the activation of the UPS^12,33,34^. Accordingly, in the next experiment, intra-PRh microinjections of the IP_3_R antagonist xestospongin C (XeC) prior to object memory reactivation blocked object memory updating (Fig. 6). A repeated measures ANOVA did not indicate a significant interaction. There were, however, main effects of drug, *F*(1,10)=5.33, *p*=.044, and context, *F*(1,10)=5.18, *p*=.046. The vehicle/same alternate context condition displayed significantly lower test phase object exploration compared to the XeC/same alternate context condition, *t*(10)=−3.18, *p*=.01. Further, the vehicle/same alternate context condition had lower test phase exploration than the vehicle/different alternate context, *t*(10)= −3.30, *p*=.008, and XeC/different alternate context, *t*(10)= −3.05, *p*=.012, conditions. There were no statistical differences between the sample phase exploration of any condition, but there was a significant main effect of context in reactivation phase exploration, *F*(1,10)=5.29, *p*=.044. However, the reactivation phase exploration means differed by a fraction of a second, so this small difference in exploration is likely not critical for test phase performance.

**Figure 6.**
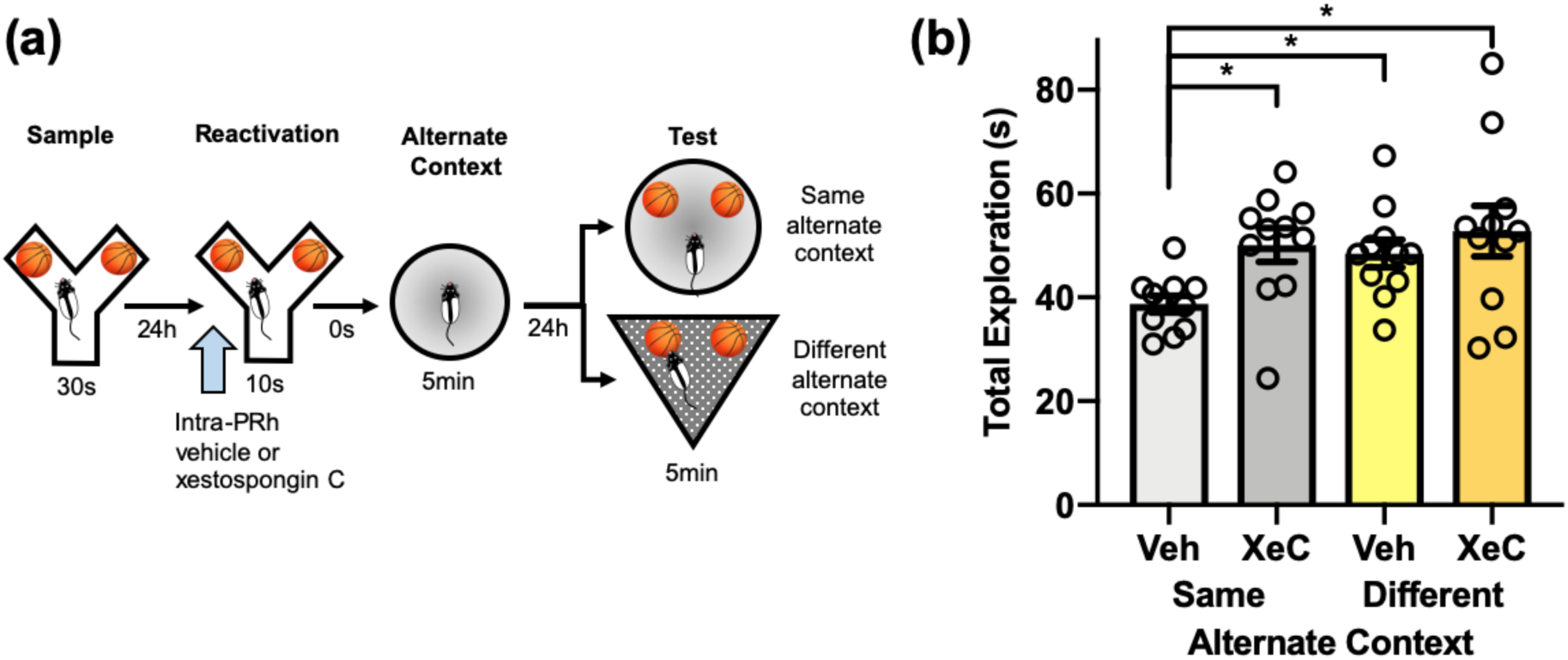
Blockade of intracellular IP_3_ receptors disrupts reactivation-based memory updating in the PROMM task. **(a)** Microinjections (blue arrow) of the IP_3_ receptor antagonist XeC or its vehicle were administered into PRh 20 min before memory reactivation in the Y-apparatus (n=11). **(b)** Rats administered pre-reactivation XeC displayed greater object exploration in the same alternate context test condition compared to rats in the vehicle/same alternate context condition. Rats in the different alternate context condition displayed greater object exploration than the vehicle/same alternate context condition, regardless of drug. \**p*<.05.

### CaMKII activity in PRh is critical for object memory updating in the PROMM task

Elevated intracellular Ca^2+^ activates CaMKII, which is involved in the phosphorylation and translocation of proteasomes to the synapse^12,33,34^. To investigate whether CaMKII might be involved in connecting M_1_ mAChR activation to UPS activity for reactivation-induced synaptic remodelling, we assessed whether inhibition of CaMKII during reactivation would prevent memory updating observed in the PROMM task. Intra-PRh microinjections of the CaMKII inhibitor KN-93 blocked object memory updating (Fig. 7). A repeated measures ANOVA indicated a significant interaction, *F*(1,11)=5.19, *p*=.044, and a main effect of context, *F*(1,11)=17.67, *p*=.001. Rats in the vehicle/same alternate context condition had lower exploration than the KN-93/same alternate context condition, *t*(11)=-2.86, *p*=.015. In addition, rats in the vehicle/same alternate context condition had lower test phase object exploration than rats in the vehicle/different alternate context condition, *t*(11)=-3.90, *p*=.002, and KN-93/different alternate context condition, *t*(11)=-4.46, *p*=.001.

**Figure 7.**
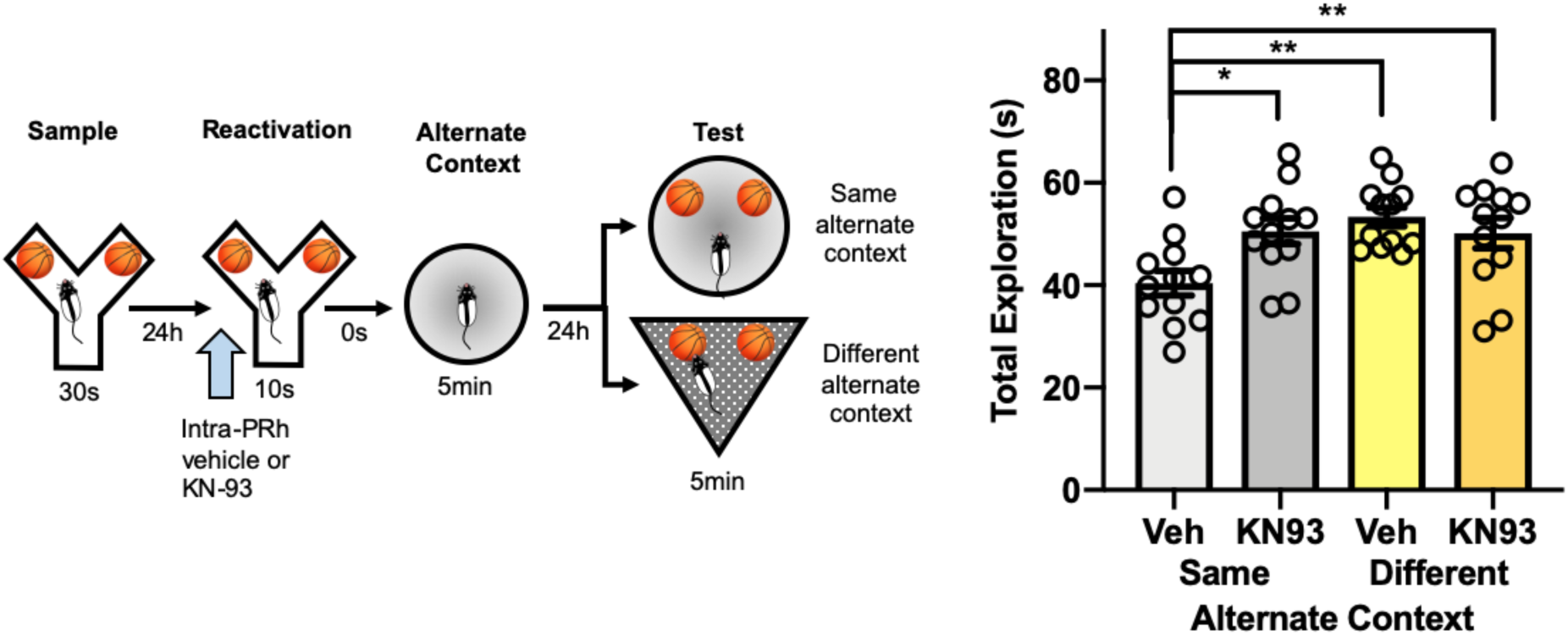
Inhibition of intracellular CaMKII within PRh blocks apparent memory updating in the PROMM task. **(a)** Microinjections (blue arrow) of KN-93, a CaMKII inhibitor, or its vehicle were administered into PRh immediately prior to memory reactivation in the Y-apparatus (n=12). **(b)** Rats in the KN-93/same alternate context condition had greater object exploration in the test phase compared to rats in the vehicle/same alternate context condition. Rats in the different alternate context had elevated test phase object exploration, regardless of drug condition. \**p*<.05, \*\**p*<.01.

## Discussion

Here we present a novel rodent paradigm that can be used to assess post-reactivation, qualitative updating of previously established object memories. Using the PROMM task, we demonstrate that novel contextual information appears to become incorporated into an existing object memory when presented following reactivation of that memory. This memory modification is constructive and reactivation- and time-dependent, and requires cholinergic signaling and UPS activation within PRh. To our knowledge, this is the first direct demonstration of cholinergic system involvement in this form of memory updating.

While our group has previously implicated ACh and downstream signaling molecules in destabilization of object memories^29,30^, designing a new reconsolidation-mediated memory updating task for rodents was a critical next step to address the involvement of this mechanism in memory updating, specifically. Tasks illustrating retrieval-related amnesia in rodents have contributed significantly to the study of neural mechanisms underlying reconsolidation, and this kind of memory modification likely has relevance to understanding long-term memory interference; however, weakening or ‘erasure’ represent only a sub-set of memory changes. Other forms of memory updating likely involve reactivation-induced integration of relevant information to maintain the adaptiveness of the memory network^6^. Indeed, there is evidence of reactivation-dependent, time-sensitive, and constructive declarative memory updating in humans^28^. The present results are highly suggestive of similar memory modification in rodents, and the PROMM task should prove valuable in continuing to uncover the neurobiological mechanisms responsible for this important cognitive function.

We developed the PROMM task to demonstrate information integration into a consolidated memory trace within the transient post-reactivation reconsolidation ‘window’ in rodents. Whereas other recently published memory modification tasks have produced informative results by presenting ‘updating’ information simultaneously with a reminder cue at the time of reactivation^25–27^, the present study appears to be the first to demonstrate constructive memory modification during the post-reactivation period. A distinct reactivation session, absent of explicit updating information, is a noteworthy adjustment compared to past memory updating tasks, as this procedure enables analysis of reactivation/destabilization mechanisms separately from mechanisms for processing the updating information. Moreover, the current design is better suited to address a central tenet of reconsolidation theory, which specifies a time-dependent window of lability – and hence modifiability – *following* memory reactivation and is also less likely to support alternative explanations in terms of novel learning episodes merely producing new memories that compete with the original memory^35^.

Similar to the methodology used by Hupbach and colleagues (2007)^28^ in humans, the current results appear to demonstrate constructive memory updating in a rodent model. Importantly, in the PROMM task, rats displayed familiarity for the sample object-context configuration during the test phase, indicating that the original memory trace remained intact after updating the memory trace with new contextual information. Correspondingly, Kwapis et al. (2019)^27^ recently developed a reconsolidation-based object-location memory updating task, demonstrating that both the original object location and an updated object location were treated as familiar compared to a novel object location. Together, the Kwapis et al. (2019)^27^ findings and the current results bode well for use of such rodent models to study the neural mechanisms of long-term memory modification and information integration. Interestingly, in the same study, Kwapis and colleagues^27^ reported deficits in object-location memory updating in aged mice; this could be explained by age-related decline in cholinergic system functioning^36^, in line with our present findings that ACh is important for reconsolidation-mediated memory updating.

The present data are consistent with the notion that the consolidated object memory is updated with the alternate context information presented during the post-reactivation period. Such a process would seem to require the object memory to enter a labile state via memory destabilization. Previously, activation of the UPS, which is involved in degradation of postsynaptic proteins, has been implicated in destabilization of various forms of memory^9,14,30,37,38^ and has been implicated in apparent memory updating^9,26^. Thus, the current results with proteasome inhibition by ß-lactone, which blocked the object memory updating effect in the PROMM task, is consistent with this speculated mechanism of memory modification that requires reactivation-induced destabilization of the original memory trace. However, as UPS activity has also been implicated in memory acquisition^33^, additional work will be required to clarify the role of the UPS in the present results.

Consistent with our past work on object memory destabilization, the findings of the current study suggest involvement of an upstream mAChR signaling pathway in recruitment of the UPS for memory trace modification. We previously linked M_1_ mAChR activation to IP_3_R stimulation and subsequent UPS activity for memory destabilization in an object recognition reconsolidation task^29,30^. We proposed that the M_1_ mAChR second messenger IP_3_ initiates mobilization of intracellular Ca^2+^ stores from the endoplasmic reticulum^32^, and this increase in cytosolic Ca^2+^ could promote downstream cellular responses to propagate protein degradation. Increases in Ca^2+^ recruit CaMKII, which has been found to activate and translocate proteasomes to dendritic spines, likely to degrade synaptic proteins involved in memory trace stabilization^12,34,39^. Accordingly, the present results implicate all of the previously identified components of this pathway for destabilization in object memory updating and extend these findings to include a role for CaMKII in this process, providing a viable signaling link between M_1_ and IP_3_ receptor activation and stimulation of the UPS. Thus, the results from the present study strongly suggest that an M_1_ mAChR-initiated mechanism for memory destabilization is necessary for qualitative modification of cortical memory representations.

Thus, ACh appears to act as a neuromodulator of memory updating, but it is likely not the only classical neurotransmitter to be involved in memory trace remodelling. Prediction error and other forms of salient novelty at the time of memory reactivation are strongly implicated in memory destabilization under certain conditions^40,41^, perhaps to signal the opportunity for memory updating. In addition to past work implicating ACh in novelty-regulated memory destabilization, there appears to be a strong role for dopamine in responding to prediction error in various reconsolidation paradigms^42–45^. Thus, further investigation is required to delineate the specific roles of various neurotransmitter systems in memory destabilization and modification; the PROMM task should prove highly valuable for such studies.

In conclusion, here we present a novel behavioural paradigm that should greatly facilitate the study of the dynamics of long-term memory storage and modification. As this task is based on the object recognition paradigm, it should produce results that could more feasibly apply to human declarative memory processes compared to similar findings that might be reported using fear conditioning or other tasks that predominate in the reconsolidation field. A rodent model with relevance to human declarative memory should prove essential for studying the basis of normal declarative memory change with time, as well as integration of inaccuracies and irrelevant information, processes that likely contribute to development of false and interfering memories. The present study provides a significant first foray into the analysis of the neurobiological basis of memory updating by providing compelling evidence for an important role of cortical cholinergic transmission in integrative object memory updating. This, to our knowledge, represents the first direct demonstration of this cognitive function for ACh, which has long been implicated in other aspects of learning and memory. Should this mechanism generalize to other forms of memory, such as fear conditioning, it could open important avenues for potential treatment of human disorders characterized by intrusive maladaptive memories, as cholinergic drugs are well established and tolerated for various other human disorders. Moreover, the current findings are potentially relevant to other aspects of human cognitive dysfunction more directly related to reduction in cholinergic system efficacy, such as in Alzheimer’s disease and normal aging^46,47^; both are characterized by severe learning and memory deficits, as well as cognitive inflexibility. The current results suggest that a subtler impairment in the updating of established long-term memories, rather than an exclusive disruption of new learning, could contribute to these symptoms. The data presented here, along with a powerful new tool in the form of the PROMM task, should help to address many of these issues in the coming years.

## Methods

### Animals

Male Long Evans rats (Charles River, NY) were housed on a 12-h reverse light cycle (lights off 8:00-20:00). Behavioural testing occurred during the dark phase of the cycle. All rats were pair-housed in standard cages with minimal enrichment. Rats were handled regularly during the first week of arrival to the facility. A restricted food regimen was implemented, and behavioural testing commenced once rats reached 85% of their typical body weight (approximately 350-425g at the start of testing). Rats were allotted 20g of 18% rodent chow each day, and water was accessible *ad libidum*. All procedures involving the use of animals were approved by the Animal Care Committee of the University of Guelph and followed the guidelines of the Canadian Council of Animal Care.

### Surgery

Bilateral PRh cannula implantation surgeries occurred when the rats were 250-320g in body weight. Rats were anaesthetized with inhalant isoflurane throughout the surgery. Medicam (concentration of 5mg/kg, subcutaneous injection) and lidocaine (20mg/kg, subcutaneous injection at incision site) were administered prior to surgery. Baytril (5mg/kg; intramuscular injection) was administered at the start of surgery. Rats were positioned in a stereotaxic frame (incisor bar set to –3.3mm). A 3-4cm incision was made across the rat’s head (anterior-to-posterior), and the skin was retracted to expose the skull. Bone screws were inserted into the surface of the skull (2 posterior to coronal suture, 2 posterior to lambdoidal suture) to secure the dental acrylic that stabilizes the guide cannula. Holes were drilled into the skull at the target coordinates for guide cannula implantation. Guide cannula placement was determined according to the following coordinates^47^ in reference to bregma: anteroposterior –5.5 mm, lateral ±6.6 mm, and dorsoventral –6.5 mm. Dental acrylic was applied to the surface of the skull around the cannulas. Skin was sutured around the dried dental acrylic. Dummy cannulas were inserted into the guides, extending 1.1mm past the tip of the guide, to prevent blockage. Rats recovered in clean cages over heating pads for 1-2h following surgery. Rats recovered for 1 week before behavioural experimentation.

### Microinjection Procedure

Dummy cannulas were removed immediately before the microinjection and placed on a sterilized surface. Drug infusers were connected to the end of tygon tubing that was secured to a glass 1.0µL Hamilton syringe. Hamilton syringes were guided by a Harvard apparatus syringe pump to administer the drug through the tubing at a constant rate of 0.5µL/min for 2-min. Drug infusers remained in the guide cannula for an additional 2-min after infusion to ensure adequate diffusion of the drug away from the infuser tips. Dummy cannulas were reinserted following microinjections. Rats were habituated to the microinjection procedure with simulated microinjections (drug infusers are inserted into the guide cannula for the same duration of time as typical microinjections, but no drug was delivered) on two separate occasions the week prior to behavioural testing.

### Histology

Cannula guide placement was verified following behavioural testing (Fig. 8). Rats were anesthetized with intraperitoneal injection of 1.2mL/kg of Euthansol (concentration of 82mg/mL), and perfused transcardially with phosphate buffered saline (PBS), followed by 4% neutral buffered formalin. Brains were extracted and stored in 4% formalin in a 3°C fridge for at least 24h. Brains were transferred to a solution of 20% sucrose in PBS, and left on an orbital shaker until they sank. Brains were sliced with a cryostat to 50μm in width, and every 3^rd^ slide was mounted onto a gelatin-coated glass microscope slide and thionin stained.

**Figure 8.**
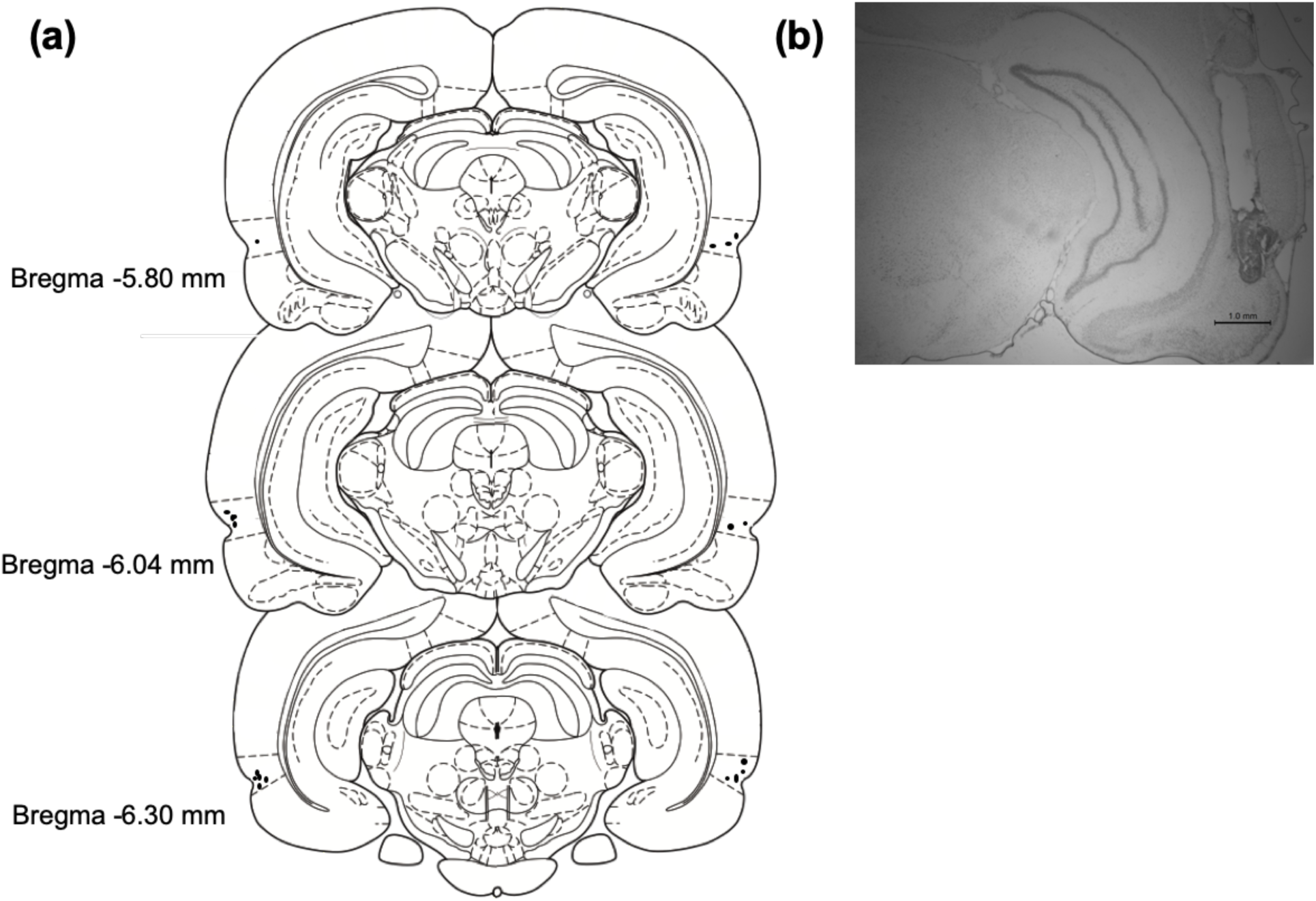
Schematic representation of cannula tip placements for microinjections targeting PRh from a representative cohort of cannulated rats. **(a)** Cannula implantation placements of cohort used for the pirenzepine experiment of this study (n=10). Dotted lines above and below the marked cannula placements (black dots) indicate the dorsal and ventral borders of PRh. **(b)** Micrograph of a typical cannula guide tract extending to PRh.

### Drugs

#### Systemic drugs

Scopolamine hydrobromide trihydrate (Sigma Aldrich), the mAChR antagonist, was dissolved in 0.9% physiological saline at a dose of 0.8mg/kg, and was administered with intraperitoneal injections 20-min prior to the reactivation phase in the object memory modification task.

#### Intracranial drugs

The M_1_ mAChR antagonist pirenzepine (Sigma Aldrich) was dissolved in 0.9% physiological saline at a concentration of 20µg/µL. The irreversible 20S proteasome inhibitor clasto-Lactacystin β-lactone (Sigma-Aldrich) was dissolved in 2% dimethyl sulfoxide (DMSO) and 1N HCl, neutralized with NaOH and concentrated with 0.9% physiological saline to 32ng/µL. Xestospongin C (Sigma-Aldrich), the IP_3_R antagonist, was dissolved in PBS and diluted to 0.3ng/µL. All intracranial drugs were microinjected immediately before the reactivation phase on a given PROMM trial, with the exception of XeC, which was administered 20-min prior to the reactivation phase. Vehicle for each experiment corresponded to the solvent used in the preparation of each respective drug.

### Statistical Analysis

Object exploration was manually scored using customized scoring software, allowing real-time scoring of each testing session. All behavioural testing was video recorded for later offline rescoring. Total object exploration (left object exploration + right object exploration) was the primary measure used in behavioural testing. An exploration bout was initiated the instant the rat’s nose was within 2cm of the object, and the bout terminated once the rat disengaged with the object. Chewing or climbing on the object were not considered object exploration. Exploration during the sample and reactivation phase of each group was analyzed to ensure that any differences in the test phase exploration were not attributable to exploration differences in the study phases of the PROMM task. The results of these analyses are only presented in cases where they were found to be significant.

Analysis of variance (ANOVA) was used to investigate statistical interactions or main effects in 2×2 (alternate context condition, drug) experimental designs. One-way ANOVA was used to evaluate group differences in the Y-apparatus control experiment. Planned comparisons established prior to performing an ANOVA were evaluated with two-tailed paired-samples or independent-samples t-tests. Bonferroni correction was used to adjust the α-level accordingly for planned comparisons and *post hocs*. Two-tailed independent-samples t-tests analyzed differences between alternate context conditions for the PROMM task and its corresponding control experiments. All t-tests in these experiments used α=0.05, excluding cases employing the Bonferroni correction. All data sets were analyzed using IBM SPSS Statistics 26 software.

### Post-reactivation object memory modification (PROMM) task

Rats were handled and habituated to all contexts involved in the PROMM task for 5 min each on two occasions the week before behavioural testing. Figure 1 illustrates the procedure for the PROMM task. The post-reactivation contexts, test phase contexts, and drug conditions were all counterbalanced across trials. Each of the three apparatuses were designated to an individual testing room to ensure that the contexts were as distinct as possible. Distinctive contextual cues, consisting of pictures of various shapes and colours, hung on the walls of the testing rooms. The white plexiglass Y-shaped apparatus was 40cm tall, and the arms were 27cm long and 10cm wide. The triangle apparatus walls were made of white corrugated plastic, with a flat black rubber floor. The posterior wall was 75cm long and other two walls were 60cm long. The walls of the triangle apparatus were 60cm tall. The circle apparatus was made of navy blue plastic, with a floor made of black fine-grain waterproof sand paper, reaching 48cm in height with a diameter of 53cm.

The PROMM task required three consecutive days of testing per trial, as it involves two 24-h delays. On the first day, the sample phase, rats explored two identical copies of a novel object for a total of 30s or a 3-min session, whichever occurred first, in the Y-shaped apparatus^49^. Twenty-four hours later, during the reactivation phase, the object memory was reactivated with a brief re-exposure (maximum 10s exploration or full 2-min session length) to the sampled objects in the same Y-shaped apparatus. Immediately following this object re-exposure, the rat was placed into an empty alternate context (either a circular apparatus or a triangular apparatus) to freely explore for a 5-min period. The goal here is to manipulate the contextual information associated with the object memory; the post-reactivation context information should integrate with the labile object memory representation, thereby producing a familiarity-like response when the object and alternate context are subsequently presented together in the test phase. During the test phase, 24h after the reactivation phase, exploration of the sampled objects was measured in either the same alternate context as the one explored post-reactivation or a different alternate context. Drug conditions were run within-subjects for all experiments except for the ß-lactone experiment; ß-lactone is irreversible, so drug was a between-subjects factor for this experiment. All rats were run on one trial per condition, except in the scopolamine and KN-93 experiments, in which all rats were run on two trials per condition to reduce variability.

Rodents preferentially explore novel stimuli more than familiar stimuli^31^, so greater test phase object exploration in one condition compared to the other was considered reflective of novelty-induced preferential exploration, and a reduction in this exploration was taken as an index of a familiarity response to the object-context configuration.

## Contributions

K.H.J. and C.E.W. contributed equally to the study; K.H.J., C.E.W., C.M., C.S., D.O., and K.A.M. performed experiments; K.H.J., C.E.W., and B.D.W. performed data analysis; K.H.J., C.E.W., and B.D.W. designed the study; K.H.J., C.E.W., and B.D.W. wrote the paper.

## Acknowledgements

Funding for this research was provided by a Natural Sciences and Engineering Research Council of Canada (NSERC) Discovery Grant to B.D.W. (400176), as well as an NSERC Canada Graduate Scholarship-Doctoral award to C.E.W. and a Canada Graduate Scholarship-Master’s award to K.H.J.

## Competing interests

The authors declare no competing interests.

